# Defective NET clearance contributes to sustained FXII activation in COVID-19-associated pulmonary thrombo-inflammation

**DOI:** 10.1101/2020.12.29.424644

**Authors:** Hanna Englert, Chandini Rangaswamy, Carsten Deppermann, Jan-Peter Sperhake, Christoph Krisp, Danny Schreier, Emma Gordon, Sandra Konrath, Munif Haddad, Giordano Pula, Reiner K. Mailer, Hartmut Schlüter, Stefan Kluge, Florian Langer, Klaus Püschel, Kosta Panousis, Evi X. Stavrou, Coen Maas, Thomas Renné, Maike Frye

## Abstract

**Background:** Coagulopathy and inflammation are hallmarks of Coronavirus disease 2019 (COVID-19) and are associated with increased mortality. Clinical and experimental data have revealed a role for neutrophil extracellular traps (NETs) in COVID-19 disease. The mechanisms that drive thrombo-inflammation in COVID-19 are poorly understood.

**Methods:** We performed proteomic analysis and immunostaining of postmortem lung tissues from COVID-19 patients and patients with other lung pathologies. We further compared coagulation factor XII (FXII) and DNase activities in plasma samples from COVID-19 patients and healthy control donors and determined NET-induced Factor XIII (FXII) activation using a chromogenic substrate assay.

**Findings:** FXII expression and activity were increased in the lung parenchyma, within the pulmonary vasculature and in fibrin-rich alveolar spaces of postmortem lung tissues from COVID-19 patients. In agreement with this, plasma FXII activation (FXIIa) was increased in samples from COVID-19 patients. Furthermore, FXIIa colocalized with NETs in COVID-19 lung tissue indicating that NETs accumulation leads to FXII contact activation in COVID-19. We further showed that an accumulation of NETs is partially due to impaired NET clearance by extracellular DNases as DNase substitution improved NET dissolution and reduced FXII activation *in vitro*.

**Interpretation:** Collectively, our study supports that the NETs/FXII axis contributes to the pathogenic chain of procoagulant and proinflammatory responses in COVID-19. Targeting both, NETs and FXIIa, could provide a strategy to mitigate COVID-19-induced thrombo-inflammation.

**Funding:** This study was supported by the European Union (840189), the Werner Otto Medical Foundation Hamburg (8/95) and the German Research Foundation (FR4239/1-1, A11/SFB877, B08/SFB841 and P06/KFO306).

## Introduction

The Coronavirus disease 2019 (COVID-19) which has caused over 2.6 million deaths and has infected 121 million people since December 2019, continues to be a major health care emergency. COVID-19 is associated with coagulopathy and increased risk of arterial and venous thrombosis that significantly contributes to mortality ^1^. The incidence of thromboembolic events such as deep vein thrombosis (DVT) and *in situ* thrombotic occlusions in the lung, liver, kidney, brain and heart is high in COVID-19 patients ^2–4^. Prophylactic anticoagulation has been recommended as standard therapy in COVID-19 patients. Additionally, increased cytokine levels (IL-6, IL-10, and TNF-α) and lymphopenia are reported in severe cases suggesting a cytokine deregulation as one of the hallmarks of COVID-19 ^5, 6^.

The vascular hyperinflammatory reactions in Severe acute respiratory syndrome coronavirus type 2 (SARS-CoV-2) infection promotes a prothrombotic state by the activation of various cell types including endothelial cells, platelets, and cells of the innate immune system. In particular, extensive neutrophil infiltration has been reported in the pulmonary interstitial and alveolar spaces in autopsies of COVID-19 patients ^7, 8^. Neutrophils are essential in the rapid innate immune response to invading pathogens ^9, 10^. Upon activation, neutrophils release neutrophil extracellular traps (NETs), that promote procoagulant reactions, including platelet activation ^11^ and fibrin generation ^12, 13^. Factor XII (FXII) is the zymogen of the serine protease FXIIa that initiates the procoagulant and proinflammatory contact system and thereby triggers the intrinsic pathway of coagulation and the bradykinin-forming kallikrein kinin system, respectively (reviewed ^14^). NETs bind FXII zymogen ^13^ and induce coagulation in plasma samples in a FXII-dependent manner ^15^. NETs are cleared from tissues and the circulation by endogenous deoxyribonucleases (DNases). We previously showed that defective DNase activity augments NETs-mediated occlusive clot formation and organ damage in non-viral systemic inflammation sepsis models ^16, 17^. Recent studies have demonstrated increased levels of NET biomarkers in serum from COVID-19 patients ^18, 19^. Furthermore, the accumulation of NETs was demonstrated in fixed lung tissues, as well as in tracheal aspirates of COVID-19 patients on mechanical ventilation ^20^. Taken together, there is accumulating evidence that NETs and the coagulation system may be causally related to the pathophysiologic manifestations of COVID-19.

In the present study, we characterize a crosstalk of innate immune cells with FXII and the contact system that contributes to adverse thrombo-inflammatory reactions in COVID-19. Interference with the NETs/FXII axis offers novel therapeutic strategies for coping with thrombo-inflammation in COVID-19. We screened postmortem lung tissues from COVID-19 patients (n=3) using a systemic proteomics approach. Systems biology found that FXII expression, as well as pathways involved in complement activation, platelet activation and neutrophil degranulation are increased in COVID-19 infected lungs. We show that NETs colocalize with the active serine protease FXIIa, in COVID-19 lung tissues indicating that NETs provide a platform for FXII contact activation in COVID-19. Ultimately, we found that DNase activity and NETs degradation by DNases is impaired in plasma from COVID-19 patients (n=43) compared to plasma from age- and sex-matched healthy donors (n=39), providing a rationale for sustained NETs-induced FXII activity in COVID-19. Taken together, the NETs-FXII axis contributes to COVID-19-associated thrombo-inflammation and can provide novel therapeutic strategies for interference with virus-associated pulmonary thrombo-inflammation.

## Methods

### Acquisition of platelet poor patient and control plasma samples

Heparinized plasma samples (n=43) were obtained from SARS-CoV-2 positive patients hospitalized at the University Medical Center Hamburg, Germany. The inclusion criterion was positive identification of a SARS-CoV-2 infection via PCR analysis. Heparinized plasma samples (n=39) from healthy donors were obtained from voluntary blood donations at the Institute of Transfusion Medicine at the University Medical Center Hamburg, Germany. Heathy donor samples were age- and sex-matched to COVID-19 plasma samples. Upon collection, all heparinized plasma samples were stored at −80°C for further analysis.

### Quantification of DNase activity in plasma

DNase activity was determined using the DNase Alert QC System (AM1970, Thermo Fisher Scientific). Substrate solution and 10X nuclease buffer, both provided in the kit, were dispensed into a black 96-well plate. Heparinized plasma was diluted 20-fold in nuclease-free water and added to the respective wells. Fluorescence was measured at excitation/emission maxima of 530/580 nm in a microplate fluorometer (Tecan Spark 10M).

### Detection of DNase activity by single radial enzyme diffusion (SRED) assay

DNase activity was measured as previously described ^21^. In brief, plasma from COVID-19 patients was loaded into wells of an agarose gel that contained DNA from salmon testes (D1626, Sigma) and SYBR Safe (S33102, Invitrogen). After incubating the gels at 37°C in a humid chamber, DNA fluorescence was recorded with a fluorescence scanner (Bio-Rad ChemiDoc™ Imager). The diameter of DNase digestion was measured using ImageJ.

### NET degradation assay

Purified human neutrophils from healthy volunteers (5×10^4^ per well) were seeded in serum-free Dulbecco’s Modified Eagle’s Medium (DMEM, Life Technologies) onto 96-well microplates. Neutrophils were activated with 10 nM phorbol 12-myristate 13-acetate (PMA, Sigma Aldrich) for 4h at 37°C, with 5% CO_2_ and humidity between 85-95%. On completion of incubation with PMA, DMEM was aspirated and neutrophils were stored overnight in PBS at 4°C. The following day, serial dilutions of Pulmozyme (500 μg/ml – 7.8 μg/ml) and 10-fold diluted plasma [in HBSS (Life Technologies) with divalent cations] were added to each well and co-incubated with formed NETs for 3h at 37°C with 5% CO_2_ and humidity between 85-95%. Plasma was subsequently removed from the plates and fixed with 2% PFA in PBS for 1h. Plates were washed with PBS before they were stained with 1μM of cell-impermeable DNA dye, Sytox Green (Life Technologies). Fluorescence was quantified using a microplate fluorometer (Tecan Spark 10M); fluorescent images of nuclei and NETs were obtained with an inverted fluorescence microscope (Zeiss Axiovert 200M).

### Measurement of FXIIa generation (chromogenic assay)

Purified human neutrophils from healthy volunteers were seeded in serum-free DMEM onto sterile 96-well plates at a concentration of 10×10^5^ cells per well. NET formation was induced with 10 nM phorbol 12-myristate 13-acetate (PMA; Sigma) for 4h at 37°C, 5 % CO_2_ and humidity between 85-95%. Purified human zymogen FXII (30μg/ml, HTI, HCXII-0155) was incubated with neutrophils that were pre-treated with vehicle or DNase I (10U/ml, Dornase alpha, Roche, Germany) for 1h, in the absence or presence of the FXIIa inhibitor, hHA-infestin-4 (500μg/ml, CSL Behring). The enzymatic activity of FXIIa was measured photometrically using the chromogenic substrate S-2302 (1mM, Chromogenix) at an absorbance wavelength of 405nm.

### Autopsies

The COVID-19 autopsy cases (n=3) were part of a larger consecutive case series ^4^. The study cohort published by Edler et al. consisted of the 80 first consecutive COVID-19 deaths analyzed at the Department for Legal Medicine at the University Medical Center Hamburg, Germany, all of which were autopsied on behalf of the health authority. The mean age of the deceased was 79.2 years, and the sex ratio was 42% men and 58% women. The cohort included patients that deceased in hospitals, nursing homes and outside institutions. The patients were PCR-positive for SARS-CoV-2 in throat or lung swabs and had both deep vein thrombosis of the lower extremities and pulmonary embolism at the time of death. For control, tissue samples were used from non-COVID-19 infected patients (n=3) who expired in the intensive care unit of the University Medical Center, Hamburg, Germany. All patients were of Caucasian origin.

### Immunofluorescence staining of human lung tissue

For visualization of lung tissue, 300 μm vibratome cross sections of formalin-fixed control or COVID-19 lung tissue were used for staining, as previously shown ^22^. Briefly, sections were permeabilized using 0.5% Triton-X100 in PBS for 20min at RT followed by blocking with 3% BSA in PBSTx for 2h. Primary antibodies were incubated overnight at 4°C, washed three times with PBSTx and subsequently incubated with secondary antibodies overnight at 4°C before further washing and mounting in Dako fluorescence mounting medium. The following antibodies were used: anti-neutrophil elastase (5 μg/ml, ab21595, Abcam), anti-chromatin (anti-histone H2A/H2B/DNA-complex, 5 μg/ml, Davids Biotechnologie GmbH ^23^), anti-Collagen I (5 μg/ml, Ab34710, Abcam), anti-fibrin (5 μg/ml, MABS2155, Merck), AlexaFluor488-conjugated anti-FXIIa (10 μg/ml, 3F7, CSL Limited), anti-citrullinated histone 3 (1:500, ab5103, Abcam). Secondary antibodies conjugated to AlexaFluor-488, −594 and −647 were obtained from Jackson ImmunoResearch (all donkey, used in 1:200 dilution). Confocal tissue images represent maximum intensity projections of Z-stacks that were acquired using a Leica SP8 inverted confocal microscope with 10x HC PL APO CS, 20x HC PL APO IMM/CORR CS2 and 63x HC PL APO Oil CS2 objectives and Leica LAS-X software. For quantification of FXIIa pixel intensity in citrullinated histone 3 (NET)-positive was performed using ImageJ. NET-positive areas were selected in the citrullinated histone 3 channel. Then, the relative fluorescent units per area (mean fluorescent intensity) of the FXIIa signal within the selected NET-positive area was measured and the average mean fluorescent intensity of the background was subtracted. n=107 NET-positive areas from three COVID-19 lungs and n=35 NET-positive areas from three CTRL lungs were analyzed in three independent experiments.

### Detection of FXIIa by immunoblotting

For the detection of FXIIa, denatured plasma samples were separated by SDS-PAGE and transferred to PVDF membranes by semi-dry blotting. The membranes were blocked with 5% normal mouse serum (015-000-120, Jackson Laboratories, USA) and incubated overnight at 4°C. After blocking, the membranes were incubated with a goat anti-human FXII antibody (1:500, GAHu/FXII, Nordic-MUbio, The Netherlands) for 1h at RT, washed and subsequently incubated with a horseradish peroxidase (HRP)-coupled mouse anti-goat secondary antibody (1:5000, #18-8814-33, Rockland Immunochemicals, Inc., USA) for an additional hour. The membranes were then washed and developed with a chemiluminescent HRP substrate. Densitometric analysis with ImageJ was performed to quantify FXIIa protein levels, which were normalized to albumin signals assessed from Ponceau staining. FXII deficient plasma (#1200, Lot 6204, George King Bio-Medical, USA) was used as a negative control.

### Statistics

GraphPad Prism was used for graphic representation and statistical analysis of data. A two-tailed unpaired Student’s t-test was used to compare between two means. Pearson’s correlation was used to correlate clinical parameters. Differences were considered statistically significant when p<0.05. The experiments were not randomized. No blinding was done for quantifications.

### Ethics

SARS-CoV-2 positive patients plasma samples were obtained at the University Medical Center Hamburg, Germany (permit number #2322, Ethics Committee of the Hamburg Medical Association, Hamburg, Germany). Control plasma samples from healthy donors were obtained from voluntary blood donations at the Institute of Transfusion Medicine at the University Medical Center Hamburg, Germany. The requirement for informed consent was waived by the Central Ethics Committee Germany given the discarded nature of the blood samples. Access to the postmortem lung tissues was granted by the Ethics Committee of the Hamburg Medical Association with a positive opinion on the examination of the tissue obtained postmortem (reference number PV7311).

### Role of funding source

The funding agencies had no role in the study design, data collection, data analyses, interpretation, or writing of report.

## Results

### FXII is increased and activated in lung tissue from COVID-19 patients

To comprehensively screen for pathways that contribute to thrombo-inflammation in COVID-19 patients that succumbed to the disease, we performed differential proteomic analysis of postmortem lung tissue from COVID-19 patients and control patients with COVID-19-independent lung pathologies (n=3, each) (Figure 1, Supplementary table 1, and Supplementary figure 1a). We sought to identify differential *in situ* protein enrichment in the lungs and avoid accumulation of proteins that derive from DVT with pulmonary embolism (PE). Therefore, we dissected COVID-19 lung tissue sections under a stereomicroscope to remove any visible thrombi that could interfere with the analysis. Gene Ontology (GO) analysis identified enrichment of pathways related to (i) classical complement activation (green), (ii) activation of proangiogenic pathways (red) as well as (iii) platelet degranulation (magenta) and (iv) neutrophil activation and degranulation (blue) in lungs of COVID-19 patients (Figure 1b). Consistently, immunofluorescence staining of COVID-19 lung vibratome sections that allow for three-dimensional visualization of the tissue architecture showed adherent degranulated platelets at the wall of microvessels using a CD62P antibody (Supplementary figure 1b). These findings are consistent with described platelet activation in COVID-19 ^24, 25^.

**Figure 1.**
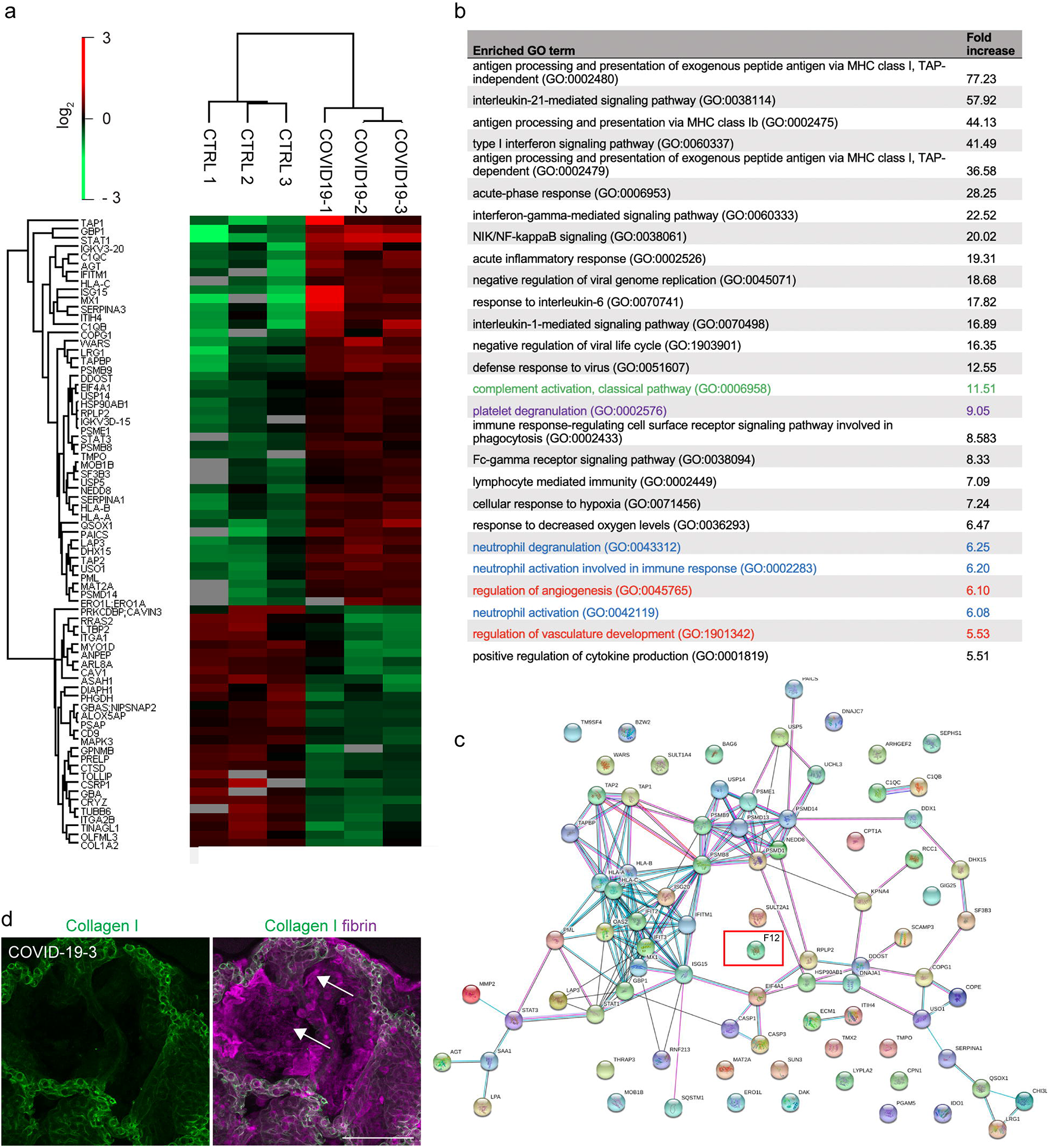
Proteomic analysis reveals enrichment of FXII and pathways involved in platelet, complement and neutrophil activation in lung tissue from COVID-19 patients. (a) Heatmap of differentially regulated proteins in the lung tissue of patients with COVID-19 and COVID-19-independent lung pathologies (CTRL) (n = 3 each). (b) Selected Gene Ontology (GO) terms of increased proteins in COVID-19 lungs highlighting pathways involved in platelet (magenta), complement (green) and neutrophil activation (blue) and angiogenesis (red). (c) Protein association network analysis (https://string-db.org) of increased proteins in COVID-19 lungs. (d) Representative immunofluorescence of lung vibratome sections using antibodies against Collagen I (green) and fibrin (magenta) shows large fibrin depositions lining the alveolar space of FXII-increased COVID-19 lungs (white arrows). Scale bar: 100 μm.

To identify key players of COVID-19-associated thrombo-inflammation in an unbiased approach, we performed protein association network analysis (https://string-db.org) of the proteomic data obtained from COVID-19 lung tissue (Figure 1c). We found peptides of the FXII sequence increased in lungs from patient COVID-19-1 and patient COVID-19-3 (Figure 1c). In line with previous studies ^2^ and indicative of a procoagulant state, proteomic analysis revealed an increase in fibrinogen protein complexes (fibrinogen alpha (log2fold change 1.73), beta (1.5), gamma chains (1.64)) in COVID-19 lungs (data not shown). Immunofluorescence analysis of COVID-19 lung vibratome sections using antibodies against fibrin and Collagen I showed dense fibrin sheet depositions at the alveolar walls (Figure 1d, arrows).

FXII is converted to FXIIa by limited proteolysis of a single peptide bond. While proteomics approach detected FXII specific peptides, the method is insensitive to discriminate whether the FXII zymogen form or the active protease FXIIa is increased in lung tissues of COVID-19 patients. To directly test if FXII is activated in COVID-19 lungs, we performed immunostaining of vibratome sections from COVID-19 lungs using antibodies against FXIIa (3F7) ^26^ and Collagen I (Figure 2a). 3F7 specifically binds to FXIIa and has no detectable cross-reactivity with the FXII zymogen form ^27^. 3F7 immunostaining demonstrated that FXII is activated in lung parenchyma, within the pulmonary vasculature and in fibrin-rich alveolar spaces of COVID-19 lung tissue (Figure 2a). In contrast, no increased FXIIa signal was detectable in lung tissue from patients that died from Acute Respiratory Distress Syndrome (ARDS) (Figure 2a). Next, we investigated if activated FXII is found intravascularly. To this end, we performed immunoblot analysis of COVID-19 plasma samples (n=19) and age- and sex-matched healthy control donor plasma (n=15) (Figure 2b+c). In agreement with our previous findings, we observed an increase in FXIIa (67%) in COVID-19 plasma versus control (Figure 2c). These results show that FXIIa is increased in lung tissue and plasma from COVID-19 patients and therefore likely contributes to pulmonary thrombo-inflammatory processes.

**Figure 2.**
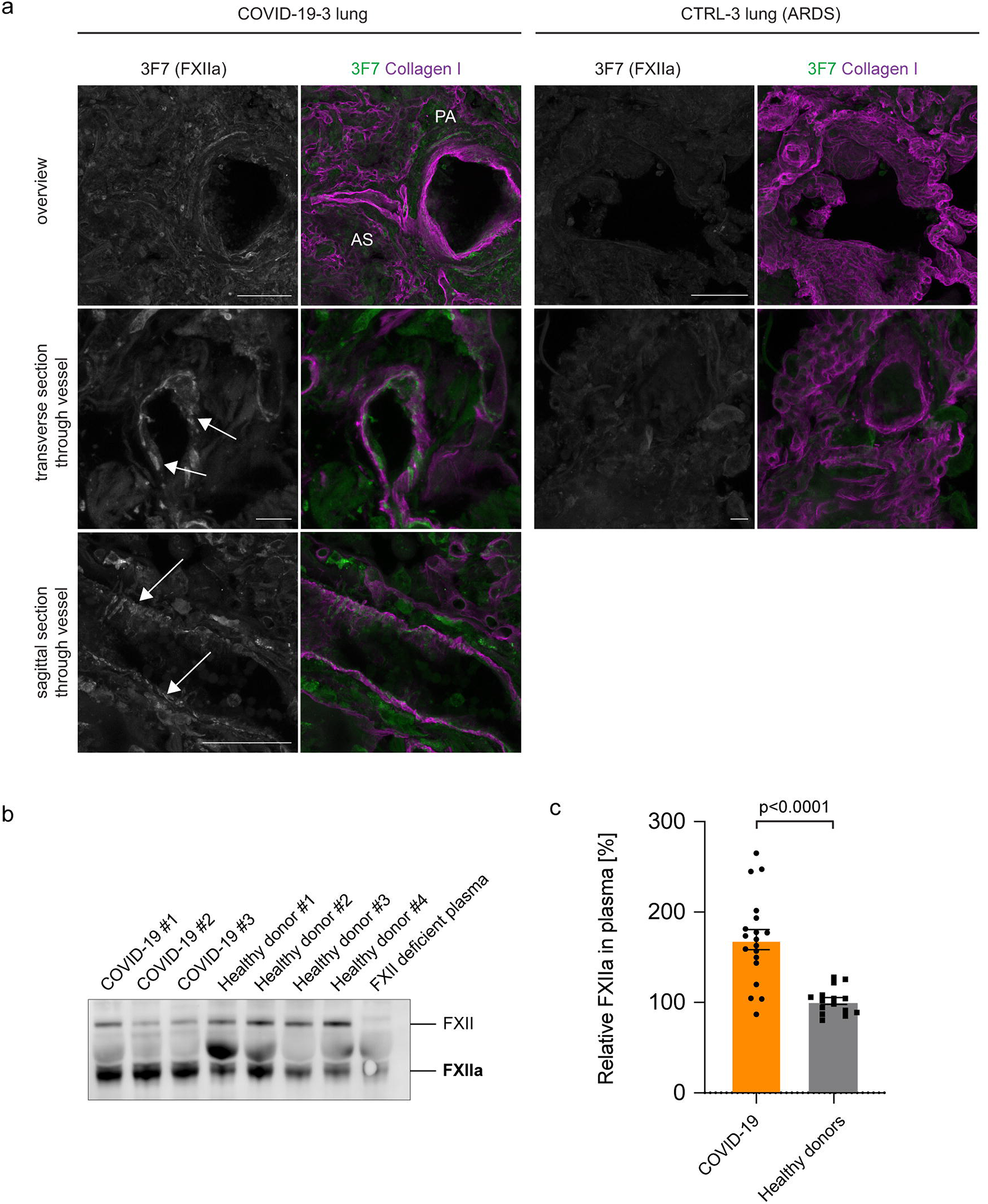
FXII is activated in COVID-19 lungs. (a) Representative immunofluorescence of vibratome sections from FXII-increased COVID-19 and Acute Respiratory Distress Syndrome (ARDS) control lungs using antibodies against FXIIa (3F7, green or grey in single channel images) and Collagen I (magenta). White arrows indicate FXIIa signal within the microvasculature. Alveolar space (AS), parenchyma (PA). (b) Representative immunoblot of COVID-19 and healthy donor plasma samples blotted for FXII (GAHu/FXII, Nordic MUbio, Netherlands). The antibody detects FXII zymogen and cleaved, activated FXII (FXIIa). As negative control FXII deficient plasma was used. (c) Quantification of FXIIa in COVID-19 plasma (n=19) and age- and sex-matched healthy donor plasma (n=15). FXIIa was normalized to plasma albumin levels. Data represent mean ± s.e.m., p-value, two-tailed unpaired Student’s t-test. Scale bar: 100 μm top/10 μm middle/50 μm bottom panel.

### NETs activate FXII in COVID-19 lungs

Next, we sought to analyse the mechanism by which FXII is activated in COVID-19. NETs have been shown to initiate FXIIa formation *ex vivo* ^13^. Furthermore, increased NET formation has been shown in serum samples and lung tissue from COVID-19 patients ^19, 20, 28^. In line with these previous findings, immunofluorescence staining of vibratome sections with antibodies against the NET biomarker neutrophil elastase (NE) and chromatin visualized degranulated neutrophils releasing elongated chromatin-positive NETs into the lung parenchyma and the alveolar spaces (Figure 3a). Consistent with the immunofluorescence, NET-specific markers such as circulating extracellular DNA and myeloperoxidase were increased in plasma samples from COVID-19 patients (n = 43) over healthy control donors (n = 39) (Supplementary table 1 and Supplementary figure 1a+b). NET biomarkers correlated with total leukocytes (r=0.80; p=0.0001), lactate dehydrogenase (cell destruction biomarker) (r=0.83; p<0.0001), and C-reactive protein (marker of inflammation) (r=0.49; p=0.0189) (Pearson’s correlation, Supplementary figure 1c-e).

**Figure 3.**
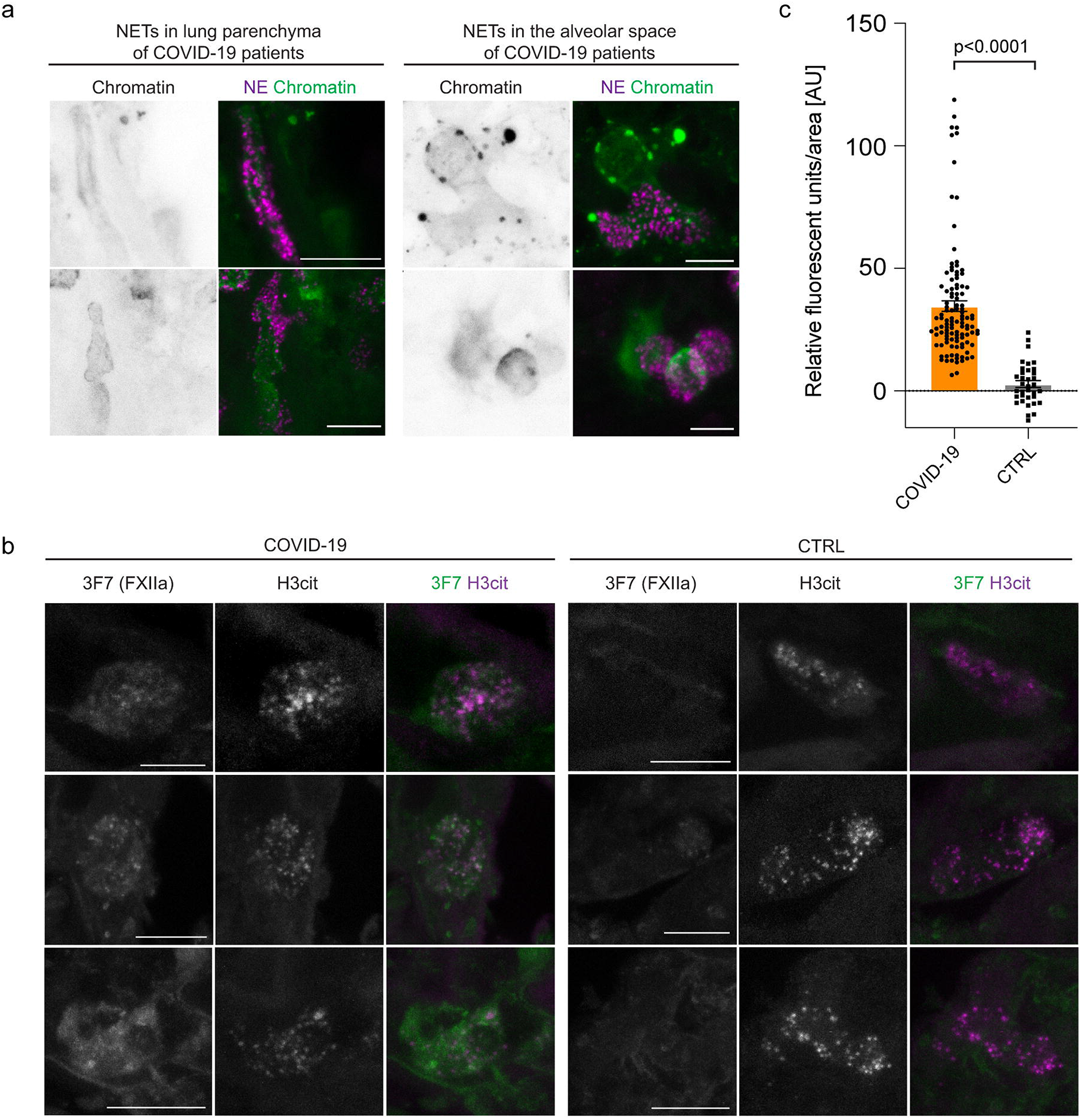
NETs and FXIIa colocalize in COVID-19 lungs. (a) High magnification immunofluorescence images of NETs in the lung parenchyma and the alveolar spaces using antibodies against NE (magenta) and chromatin (green or grey in single channel images). (b) Representative immunofluorescence of vibratome sections of COVID-19 and control (CTRL) lungs using antibodies against activated FXII (3F7, green or grey in single channel images) and histone 3 (H3cit) (magenta or grey in single channel images). (c) FXIIa pixel intensity was measured in n=107 NET-positive areas from three COVID-19 lungs and n=35 NET-positive areas from three CTRL lungs. Data represent mean ± s.e.m., p-value, two-tailed unpaired Student’s t-test. Scale bar: 10 μm (a,b).

To test if NETs can provide a scaffold to induce FXII contact activation in COVID-19 lungs, we performed high resolution immunostaining of lung vibratome sections from COVID-19 lungs using antibodies against FXIIa (3F7) and the NET component citrullinated histone 3 (Figure 3b). We quantified FXIIa pixel intensity in citrullinated histone 3 (NET)-positive and observed a 12-fold increase in colocalization of FXIIa and NET-positive structures in COVID-19 lungs compared to CTRL lungs (Figure 3c), indicating that NETs specifically induced FXII activation in COVID-19 lung tissue.

### Defective NET clearance underlies the sustained FXII activation in COVID-19 lungs

As NETs are abundant in the circulation, lung parenchyma and alveolar spaces of COVID-19 patients, we next sought to identify the underlying mechanism of NETs accumulation. Two independent experimental approaches analysing for DNase activity in plasma samples from COVID-19 patients were performed. Both assays showed that DNase activity was reduced in plasma samples from COVID-19 patients compared to levels in heathy donors (Figure 4a+b). Furthermore, defective DNase activity was corroborated in a third NETs degradation assay where *in vitro* generated NETs from healthy donors were incubated with plasma either from COVID-19 patients or healthy donors (Figure 4c). Healthy donor plasma or recombinant human DNase I (Pulmozyme, data not shown) completely degraded NETs within 3h. In contrast, COVID-19 plasma was inactive to degrade NETs (Figure 4c). We conclude that excessive NET accumulation in COVID-19 results from increased generation and impaired degradation of NETs.

**Figure 4:**
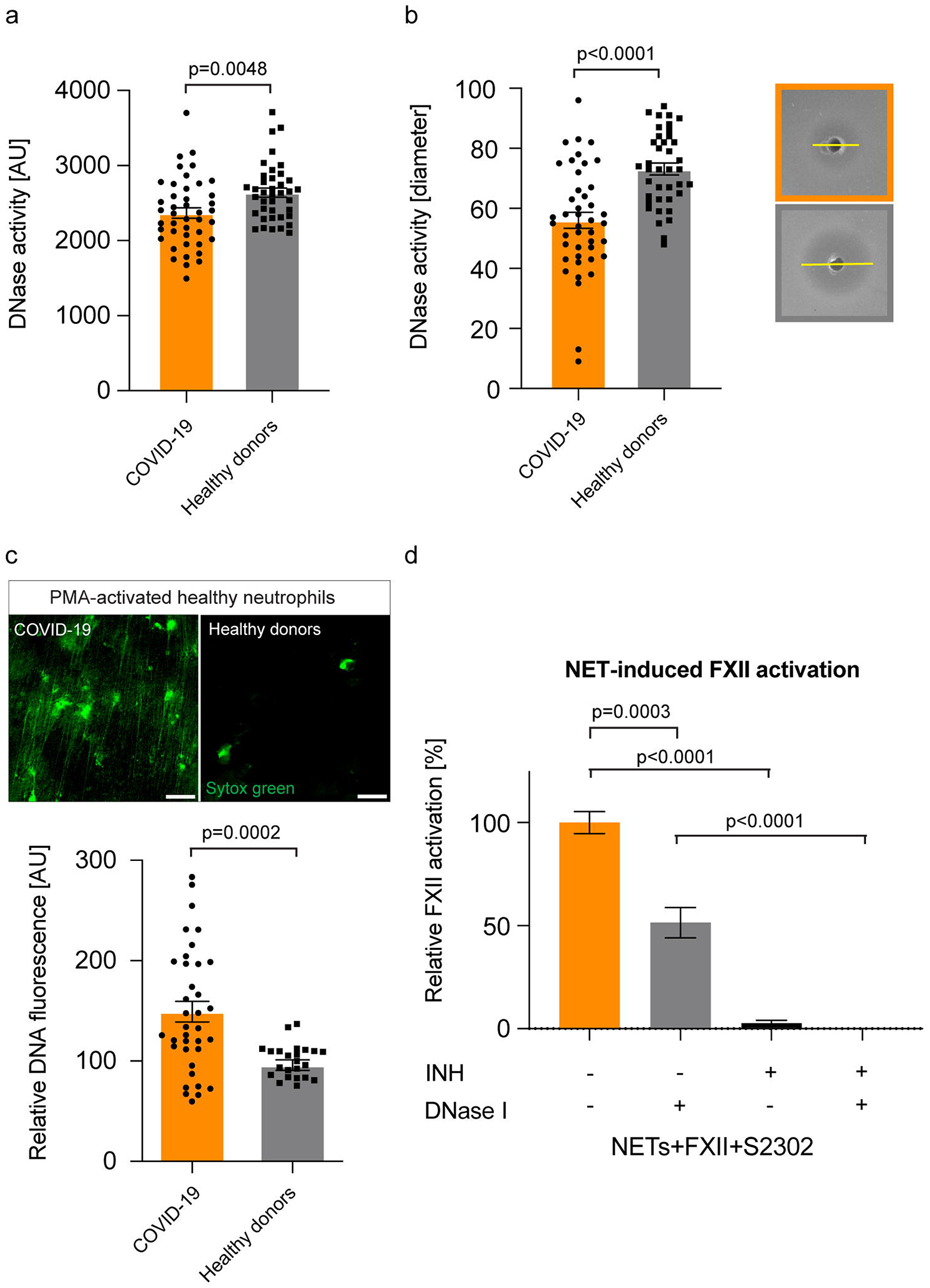
Delayed NET clearance due to defective DNase activity in COVID-19. (a) DNase activity measured in plasma samples from COVID-19 patients (n=43) or healthy donors (n=39) using the DNase Alert QC System. Data represent mean ± s.e.m., p-value, two-tailed unpaired Student’s t-test. (b) DNase activity measured in plasma samples from COVID-19 patients (n=43) or healthy donors (n=39) using the SRED assay. Representative pictures are shown for COVID-19 (orange box) and healthy donors (grey box). Data represent mean ± s.e.m., p-value, two-tailed unpaired Student’s t-test. (c) NET degradation capacity of COVID-19 plasma. DNA fluorescence was quantified from COVID-19 plasma (n=36) and healthy donor plasma (n=24). Data represent mean ± s.e.m., p-value, two-tailed unpaired Student’s t-test. Representative immunofluorescence images of *in vitro* generated NETs from healthy neutrophils are shown upon incubation with healthy donor or COVID-19 plasma. (d) Activation of purified FXII by NETs was measured by conversion of the chromogenic substrate H-D-Pro-Phe-Arg-p-nitroaniline (S-2302) at an absorption of λ=405 nm. S-2302 was added either in the presence or absence of the FXIIa inhibitor rHA-infestin-4 (INH) or DNase I (Pulmozyme). Data represent mean ± s.e.m., p-value, two-tailed unpaired Student’s t-test (n=6 replicates per time point from two independent experiments). Arbitrary units (AU), Scale bar: 25 μm (c).

To evaluate a potential therapeutic approach for interference with the NETs/FXIIa axis, we tested if accelerated NET clearance through DNase I addition impacts on FXII activation. To that end, healthy human neutrophils were activated to produce NETs (10nM, phorbol 12-myristate 13-acetate, PMA) and formation of FXIIa was analyzed by a chromogenic assay (Figure 4d). Vice versa, degradation of NETs following addition of DNase I (Pulmozyme) reduced FXIIa formation by about half (51%). The FXII inhibitor rHA-infestin-4, on the other hand, completely suppressed the FXIIa signal (Figure 4d). Taken together, we provide evidence that defective NETs clearance could underlie the sustained FXII activation in COVID-19-associated pulmonary thrombo-inflammation. Besides targeting FXIIa, these data suggest that pharmacologic dissolution of NETs can be beneficial in COVID-19.

## Discussion

Our study uncovers a role of NET-induced activation of FXII in COVID-19-associated thrombo-inflammation. Two types of thrombotic complications occur in COVID-19 patients: DVT with pulmonary embolism can be frequently observed, especially in patients receiving intensive care ^28^. On the other hand, COVID-19 patients also have a nine times higher prevalence of alveolar-capillary microthrombi compared to patients with influenza ^29^. There is a high incidence of venous thromboembolic events in severe COVID-19 cases, despite prophylactic anticoagulation ^1, 30, 31^. Even after hospital discharge of COVID-19 patients, thrombotic events can be frequently observed, demonstrating the importance of extended thromboprophylaxis ^32^. In addition to NETs/FXIIa, other procoagulant mechanisms, such as endothelial damage ^33^, hyperfibrinogenemia ^34^ and procoagulant platelets ^35^, have been shown to contribute to thrombotic processes in COVID-19. Antiviral therapies that reduce cell injury are currently limited for COVID-19 ^36^, therefore the development of new strategies that target thrombo-inflammatory mediators could contribute to mitigate COVID-19-associated thrombotic complications.

An early study showed that in-hospital mortality of patients on mechanical ventilation was reduced by 50% upon receiving therapeutic anticoagulation (AC) and a prophylactic treatment plan including a higher AC regimen could ameliorate the disease severity in COVID-19 cases ^37^. However, a recent interim statement of the ACTIV4-ACUTE study revealed that full AC was beneficial in COVID-19 patients with intermediate severity and without mechanical ventilation, but futile in COVID-19 patients with high severity on ventilation ^38^. These findings suggest that early AC treatment prevents adverse outcomes in COVID-19. Since more potent AC is often associated with an increased bleeding risk, alternative approaches, which target coagulation but spare hemostasis, are needed. Targeting the NETs/FXII axis in COVID-19 patients at an early stage of disease progression could therefore contribute to safe interference with clinical microvascular thrombosis.

COVID-19 infection has been shown to cause inflammation of endothelial cells (endotheliitis) ^29^. Proteomic analysis of COVID-19 lungs showed enrichment of pathways associated with angiogenesis and vascular development suggesting induction of blood vessel growth in COVID-19 lungs (Figure 1B). In line with this, intussusceptive angiogenesis was increased in lungs of COVID-19 patients compared to patients with influenza infection ^29^. Furthermore, we found upregulation of complement system proteins in COVID-19 lungs. This is in line with previous publications that suggested that complement and NETs contribute to COVID-19 immuno-thrombosis ^39, 40^.

Although we studied a limited number of postmortem lung tissue from COVID-19 patients, we found FXII to be increased in COVID-19 lung tissues but not in lung tissues of other lung pathologies. The 3F7 antibody, which specifically detects FXIIa ^26^, showed that expression of FXII was accompanied by an equal increase in FXII activity in lung parenchyma, within the pulmonary vasculature and in fibrin-rich alveolar spaces of COVID-19 lungs. Colocalisation of NETs and FXIIa in COVID-19 lung immunostaining indicates that NETs act as a scaffold for FXII contact activation in COVID-19 and possibly other NETs-driven disease states. We found that FXIIa is also increased in plasma from COVID-19 patients. Increased FXIIa levels in plasma together with increased endothelial barrier permeability (that leads to pulmonary edema in COVID-19 patients), suggests that leakage of FXII into subendothelial tissues might take place. In previous studies, we showed that neutrophils express a pool of FXII that is functionally distinct from hepatic-derived FXII and contributes to NETs formation ^41^, suggesting that increased FXII might additionally originate from infiltrated neutrophils in COVID-19 lungs. In line with our hypothesis that the NETs/FXII axis plays a role in COVID-19, others have shown that NETs have the capacity for initiating FXII activation ^15^. Additionally, FXII-independent pathways have been described to trigger NET-induced thrombosis ^19^.

In line with published studies ^18–20^, we showed excessive NETs accumulation in COVID-19 plasma and lung tissues. Physiologically, NET degradation is ensured by host DNases which regulate NET turnover to prevent their accumulation and downstream thromboinflammatory sequelae ^21^. Indeed, NET accumulation has been associated with reduced DNase activity in thrombotic microangiopathies and sepsis ^42^. Although the initial trigger of NET formation during COVID-19 is not fully understood, we observed reduced DNase activity in COVID-19 patient plasma which was accompanied by an impaired capacity to digest *in vitro* generated NETs from healthy neutrophils. Furthermore, improved NET clearance through exogenous DNase I reduced FXII activation *in vitro*, suggesting that COVID-19 patients could benefit from DNase I treatment. Currently, there are nine interventional clinical trials ongoing that investigate the safety and efficacy of DNases in COVID-19 patients, including recombinant human DNase I [Pulmozyme; ClinicalTrials.gov identifiers: NCT04355364, NCT04402970, NCT04445285, NCT04432987, NCT04359654, NCT04402944, NCT04524962, NCT04541979 and NCT04409925].

Besides its well-known selective role in thrombosis ^14^, FXII drives production of the inflammatory mediator bradykinin. The activity of bradykinin is controlled through degradation by several broad-spectrum cell-surface associated peptidases, including angiotensin-converting enzyme (ACE), dipeptidyl peptidase 4 (DPP4), and aminopeptidase P ^43^. Furthermore, its derivative des-Arg^9^-bradykinin is a target for degradation by ACE-2, the receptor that SARS-CoV-2 uses to enter cells ^44^. The exact role of ACE-2 remains to be shown as a recent systems biology study surprisingly showed that ACE-2 levels, which were suspected to be downregulated as a result of virus entry, were elevated in plasma samples of male COVID-19 patients ^45^. On the other hand, plasma levels of DPP4 (an uptake receptor for coronaviruses, but not for SARS-CoV-2) were reduced ^45^, suggesting dysregulated bradykinin signaling in COVID-19. Targeting bradykinin production or signaling appears to attenuate pulmonary edema in COVID-19 patients ^46, 47^. Our study provides additional evidence for a role of the kallikrein kinin-system in COVID-19; carboxypeptidase N (CPN1) was exclusively found in COVID-19 lung tissue (Figure 1c), which would result in accelerated conversion of bradykinin to des-Arg^9^-bradykinin. Finally, we found reduced levels of aminopeptidase N (ANPEP) in lungs of COVID-19 patients (Figure 1a). This enzyme (another receptor for coronaviridae, https://www.ncbi.nlm.nih.gov/gene/290) is expressed by numerous cell types, including inflammatory cells ^48^. ANPEP plays a role in the activation of chemokines and trimming of antigens prior to their presentation. Interestingly, the activity of this enzyme is competitively blocked by bradykinin (and therapeutic analogs ^49, 50^. This suggests that besides its direct proinflammatory roles, increased bradykinin levels may indirectly contribute to inflammation by repressing ANPEP activity in COVID-19 patients.

Activated FXII drives coagulation in various vascular beds ^51^ and the formation of bradykinin suggests that activated FXII itself is an attractive drug target for interference with thrombo-inflammation. Together with CSL Behring, we have previously demonstrated that targeting FXIIa with the neutralizing antibody 3F7 provides thrombo-protection and impedes bradykinin production in a preclinical setting ^27^. Recently, CSL Behring has initiated a clinical trial to test efficacy of the FXIIa neutralizing antibody CSL312 (Garadacimab, a successor molecule of 3F7) in COVID-19 patients (ClinicalTrials.gov Identifier: NCT04409509).

In conclusion, we uncovered a role of defective NETs clearance in sustained FXII contact activation in COVID-19-associated pulmonary thrombo-inflammation. Our data show that the NETs/FXII axis contributes to COVID-19-induced lung disease and suggest that therapeutic strategies targeting both NETs and FXIIa, could interfere with COVID-19-induced thrombo-inflammation and beyond.

## Contributors

H.E., C.R., C.D., and S.Ko. performed plasma experiments, M.F., JP.S., C.K., and D.S. performed lung tissue experiments. E.G., G.P., R.M., F.L. E.S., and C.M. conceived and discussed data. M.H., S.Kl. and K.Pü. provided essential resources. K.Pa. and H.S. provided essential tools. H.E., C.R., C.D., T.R. and M.F. wrote the manuscript. M.F. conceived and directed the study. All authors discussed the results and approved the final version of the manuscript.

## Declaration of interests

H.E. has a patent WO 2019/036719A2, licensed to Neutrolis Therapeutics, outside the submitted work. C.R. has a patent EP3570667A1, licensed to Neutrolis Therapeutics, outside the submitted work. K.Pa. is an employee and shareholder of CSL Limited and has patents 9.856.326, 9.856.325 and 9.518.127 issued, outside the submitted work. C.M. is part-time employee of TargED Biopharmaceuticals and has a patent WO2019185723A1 pending, both outside the submitted work. T.R. and the University of Hamburg have a patent (EU and U.S. reference numbers EP18736859.2 and US16/622064) licensed, outside of the submitted work.

## Acknowledgements

This study was supported by the European Union’s Horizon 2020 Research and Innovation Programme under the Marie Skłodowska-Curie Grant Agreement No. 840189, the Werner Otto Medical Foundation Hamburg (8/95) and German Research Foundation (DFG) grant FR 4239/1-1 (to M.F.) and grants A11/SFB877, B08/SFB841 and P06/KFO306 of the DFG (to T.R.). We thank the Institute of Transfusion Medicine and the Microscopy Imaging Facility at the University Medical Center Hamburg, Germany, for obtaining healthy control plasma samples and for technical microscopy support, respectively.

## Data sharing

All relevant data are available from the corresponding author upon request. The mass spectrometry proteomics data have been deposited to the ProteomeXchange Consortium ^52^ via the PRIDE ^53^ partner repository with the dataset identifier PXD020216.

