## Supplementary figures and images for "Defective NET clearance contributes to sustained FXII activation in COVID-19-associated pulmonary thrombo-inflammation"

### Supplementary figure 1

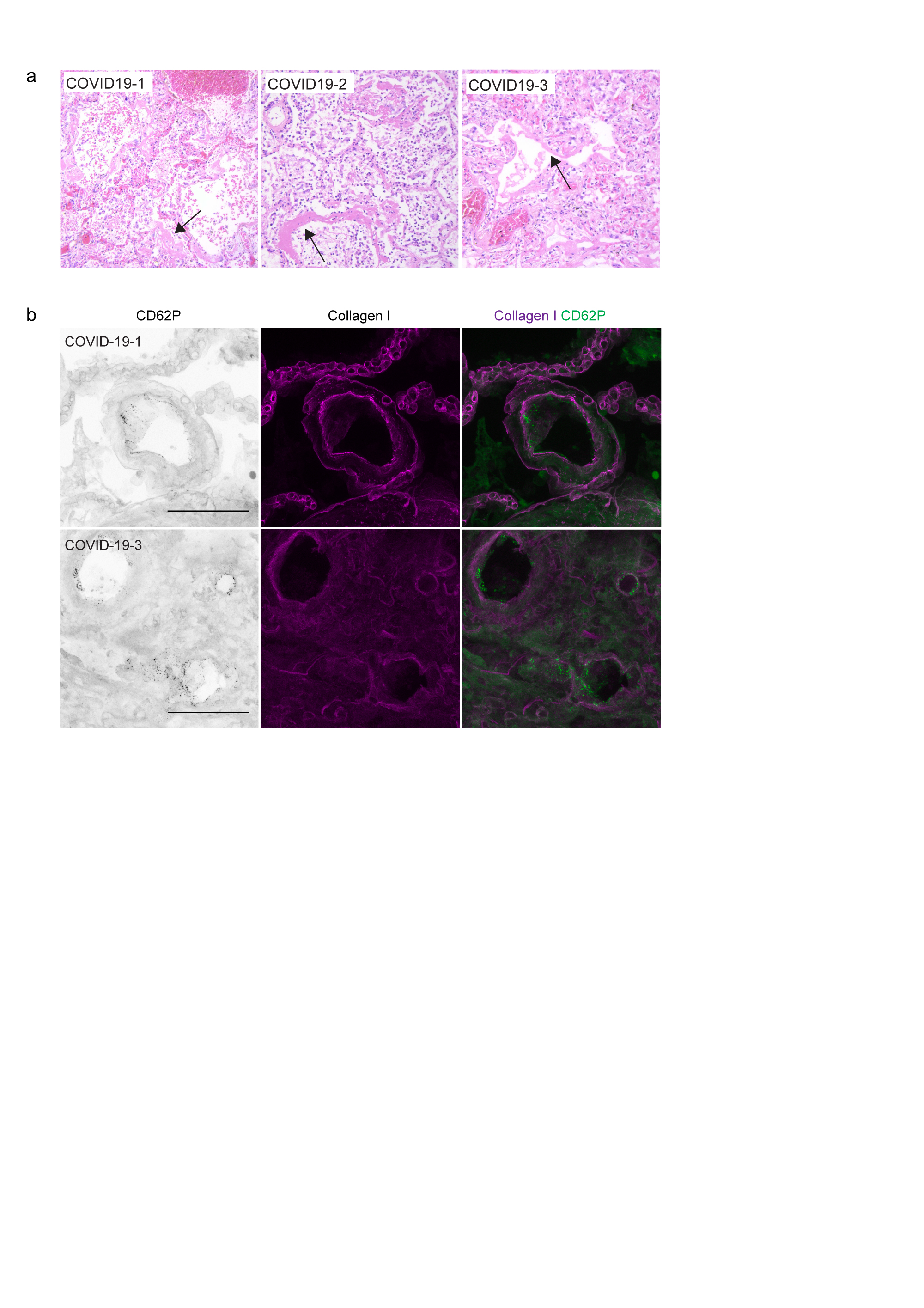

### Supplementary figure 2

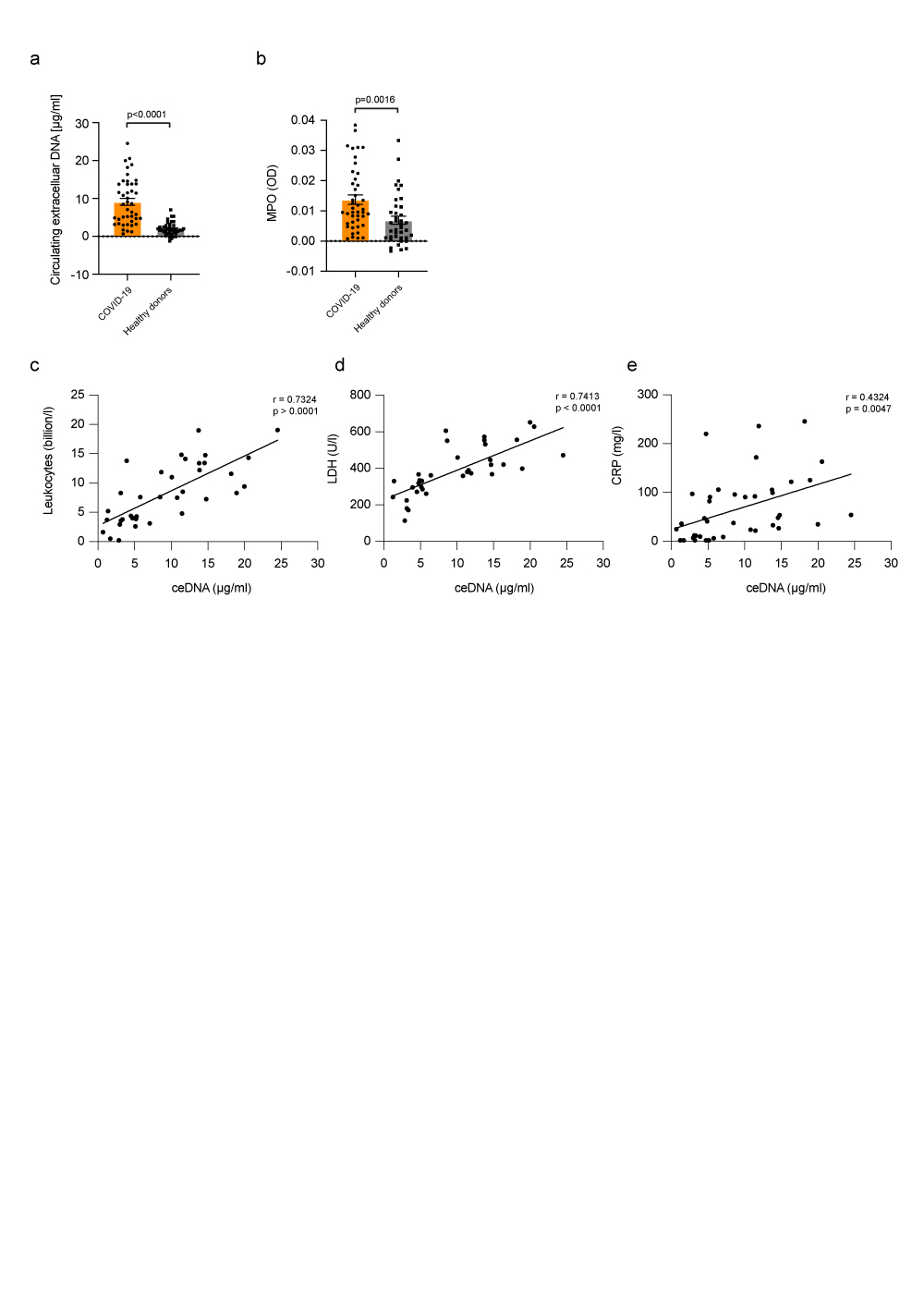

### Supplementary table 1

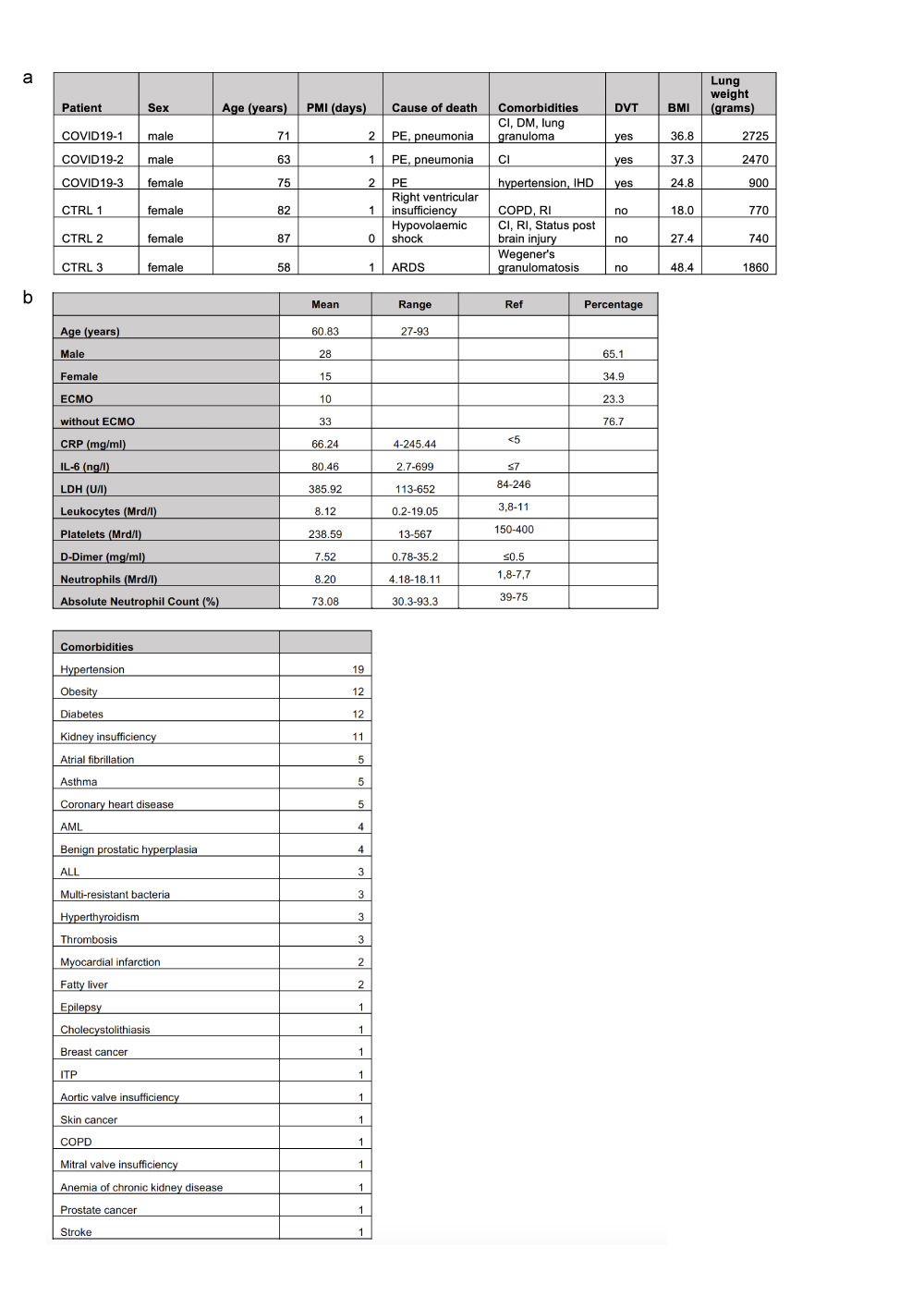
