## Supplementary data and figure legends for "Defective NET clearance contributes to sustained FXII activation in COVID-19-associated pulmonary thrombo-inflammation"

### Supplemental methods

**Sample preparation for liquid chromatography with tandem mass spectrometry (LC-MS/MS) measurements.** 4% PFA-fixed 100µm lung vibratome sections were dissected under a stereomicroscope to exclude large pulmonary embolism (PE)-derived thrombi within the tissue for subsequent LC-MS/MS measurements. Section fragments were lysed in 100 mM ammonium bicarbonate and 1% w/w sodium deoxycholate buffer boiled at 95 °C for 60 min and sonicated with a probe sonicator to destroy DNA/RNA. Plasma samples were diluted in 100 mM ammonium bicarbonate (TEAB) and 1% w/v sodium deoxycholate (SDC) buffer boiled at 95 °C for 10 min. The protein concentrations were estimated using the BCA Protein Assay kit (Pierce, Thermo Fisher) and 20 µg of protein per sample were reduced in the presence of 10 mM dithiothreitol (DTT) at 60°C for 30 min and alkylated in 20 mM iodoacetamide (IAA) for 30 min in the dark at 37 °C. Trypsin (sequencing grade, Promega) was added at a 1:100 ration (enzyme to protein) and digestion was performed overnight at 37 °C. To stop the reaction and precipitate SDC, formic acid (FA) was added to 1 % final concentration. The samples were centrifuged for 5 min at 16.000 g, the supernatant was transferred into a new tube and was dried in a vacuum centrifuge. For LC-MS/MS analyses, samples were resuspended in 0.1 % FA at a concentration of 1 µg/µl.

**LC-MS/MS measurements**. LC-MS/MS measurements were carried out on a Quadrupole Orbitrap Iontrap tribrid mass spectrometer (Fusion, Thermo Fisher) coupled to an UPLC system (Dionex Ultimate 3000, Thermo Fisher). For analysis, 1 µg of peptides were loaded by autosampler injection onto a C18 reversed phase (RP) trap column (Symmetry C18 trap column, 100 Å pore size, 5 µm particle diameters, 180 µm x 20 mm) and separated on a 25 cm C18 RP (Peptide BEH C18 column, 130 Å pore size, 1.7 µm particle diameters, 75 µm x 250 mm). Trapping was done for 5 min at a flow rate of 5 µl/min with 100 % solvent A (0.1 % FA). Separation and elution of peptides was achieved by a linear gradient from 1 to 30 % solvent B (0.1 % FA in ACN) for 60 min for the lung tissue samples and 120 min for the plasma samples.
The eluting peptides were transferred via electrospray ionization (ESI) to the Fusion mass spectrometer. MS1 scans were performed in positive mode over a scan range of 400-1300 m/z. The Orbitrap resolution was set to 120.000 with an AGC target of 2x105 and a maximum injection time of 120 ms. Peptides with charge states between 2+ - 5+ above an intensity threshold of 200.000 were isolated with a 1.6 m/z isolation window in top speed mode and fragmented with a normalized collision energy of 30 %. The fragments were measured with an Orbitrap resolution of 15,000, AGC target of 1x105 and 60 ms maximum injection time. Already fragmented peptides were excluded for 15 s.

**LC-MS/MS Data Processing.** The collected raw files were searched against the reviewed human protein database downloaded from Uniprot (April 2020 with 20365 entries) and the SARS-CoV-2 protein database (June 2020 with 14 entries) processed with the Sequest Algorithm included in the Proteome Discoverer Software (Thermo Fisher, Version 2.4.1.15). All samples were handled as individual experiments. Label-free quantification option with precursor mass recalibration and feature mapping across all samples was used. Trypsin was selected as the enzyme used to generate peptides, allowing a maximum of two missed cleavages. Oxidation of methionine, acetylation of protein N-termini and the conversion of glutamine to pyro-glutamic acid was set as variable modification. The carbamidomethylation of cysteines was selected as fixed modification. The error tolerance for the precursor search was set to 10 ppm. The fragment error tolerance was set to 0.2 Da. False discovery rate for the peptide level was set to 1 %. For quantification all identified razor and unique peptides above the FDR cutoff were considered.
The Protein areas (summed peptide areas per protein exported from Proteome Discoverer 2.4) were loaded into Perseus software (Max Plank Institute for Biochemistry, Version 1.5.8.5). The protein area values for all master proteins were used as main columns, transformed to log2 values, filtered for proteins with at least two out of three valid values in at least one group (Covid-19 or control) in the lung samples and in at least about 60 % of samples in at least one group for plasma samples. Data were then normalized by the column median. Student's t-test were calculated; hierarchical clustering was performed for proteins considered significantly different between the groups (p-value > 0.05 and log2 fold change either > 1 or < -1).

**Quantification of circulating free DNA in plasma**. Heparinized plasma was 20-fold diluted in dilution buffer containing 0.2% BSA and 4 mM EDTA in phosphate-buffered saline (PBS). Equal amounts of diluted plasma and PBS containing 1 µM of the fluorescent DNA-intercalating dye Sytox Green (Invitrogen) were mixed. Fluorescence was measured in a microplate fluorometer (Tecan Spark 10M). DNA concentrations were calculated based on a lambda DNA standard curve (Invitrogen).

**Myeloperoxidase concentration in plasma.** Myeloperoxidase (MPO) concentration was quantified in duplicates from heparinized plasma samples using a commercial human myeloperoxidase enzyme-linked immunosorbent assay kit (Duoset R&D Systems, DY3174). MPO concentration was determined according to the manufacturer's protocol. In brief, a 96-well microplate was coated with capture antibody overnight, washed and consecutively blocked for 1 h at RT. The plate was incubated with the 20-fold diluted samples and standards for 2 h at RT. Wells were washed and then incubated for 2 h with detection antibody. Washing was repeated before the streptavidin-HRP working solution was added to the wells for 20 min in the dark. Following additional washing steps, HRP-activity was detected using ABTS solution (Life Technologies). The OD was determined using a microplate reader (Tecan Spark 10M) set at 450 nm.

### Supplemental table and figures

**Supplementary table 1: Clinical data and comorbidities of COVID-19 patients in this study.**
(a) Demographics of patients used for lung tissue analysis. Lung tissue was analyzed from 3 COVID-19 patients and 3 age- and postmortem interval (PMI)- matched control patients with other lung pathologies. (b) Patient data from plasma analysis. Plasma was analyzed from 43 COVID-19 patients and extracorporeal membrane oxygenation (ECMO) status, mean clinical chemistry parameters and comorbidities are shown. Reference values (Ref) of healthy donors are included for comparison. Pulmonary embolism (PE), Acute respiratory distress syndrome (ARDS) Cardiac insufficiency (CI), Diabetes mellitus (DM), Ischemic heart disease (IHD), Chronic obstructive pulmonary disease (COPD), Renal insufficiency (RI).

**Supplementary figure 1: Characterization of COVID-19 lungs.**
(a) Histological analysis of H&E stained lung paraffin section of COVID-19 lung tissue (n=3). Hyaline membranes are indicated with black arrows. (b) Degranulated platelets adhere at microvascular vessel walls in COVID-19 lungs. Immunofluorescence of lung vibratome sections from COVID-19 patients using antibodies against Collagen I (magenta) and CD62P (green, marker for degranulated platelets). Scale bar: 100 µm.

**Supplementary figure 2. Intravascular NETs formation in COVID-19 patients**
(a-b) Circulating extracellular DNA (a) and myeloperoxidase (b) levels were measured in plasma samples from COVID-19 patients (n = 43) and healthy donors (n = 39). Data represent mean ± s.e.m. p value, unpaired Student's t-test. (c-e) Circulating extracellular DNA levels in COVID-19 plasma samples were correlated with the laboratory markers absolute leukocyte count (n = 36) (c), lactate dehydrogenase (LDH) (n = 36) (d) and c-reactive protein (CRP) (n = 41) €. Correlations with laboratory parameters were performed, when available. Linear regression, Pearson's correlation coefficients and p values are shown in each panel.
